# Apoptosis is a generator of Wnt-dependent regeneration and homeostatic cell renewal in the ascidian *Ciona*

**DOI:** 10.1101/2020.12.18.423505

**Authors:** William R. Jeffery, Spela Goricki

## Abstract

Body regeneration is unilateral in the ascidian *Ciona intestinalis*: severed basal body parts can regenerate distal structures, such as the siphons and neural complex, but severed distal body parts do not replace basal structures. Regeneration involves the activity of adult stem cells in vasculature of the branchial sac, which are induced to proliferate and produce migratory progenitor cells for the replacement of missing tissues and organs. Branchial sac-derived stem cells also replenish continuously recycling cells lining the pharyngeal fissures during homeostatic growth. Apoptosis at injury sites is an early and transient event of regeneration and occurs continuously in the pharyngeal fissures during homeostatic growth. Treatment of amputated animals with caspase 1 inhibitor, caspase 3 inhibitor, or the pan-caspase inhibitor Z-VAD-FMK blocked apoptosis, prevented regeneration, and suppressed the growth and function of the branchial sac. A pharmacological screen and inhibitory siRNA treatments indicated that regeneration and homeostatic growth require canonical Wnt signaling. Furthermore, exogenously supplied recombinant Wnt3a protein rescued both caspase-blocked regeneration and normal branchial sac growth. As determined by EdU pulse-chase studies, inhibition of apoptosis did not affect branchial sac stem cell proliferation but instead prevented the survival of progenitor cells. After bisection across the mid-body, apoptosis at the injury site occurred in the regenerating basal fragments, but not in the non-regenerating distal fragments, although both fragments contain a large portion of the branchial sac, suggesting that apoptosis is unilateral at the wound site and the presence of branchial sac stem cells is insufficient for regeneration. The results show that apoptosis-dependent Wnt signaling mediates regeneration and homeostatic growth by promoting progenitor cell survival in *Ciona*.

## Introduction

Ascidians are well known for their inability to replace the function of ablated embryonic cells and thus are often described as exemplary models for determinate or mosaic development (Jeffery, 2001; Nishida, 2004). After metamorphosis, however, ascidian juveniles and adults show impressive capacities for regeneration of injured body parts. Colonial ascidians, such as *Botryllus*, can reform the entire zooid body, including all somatic and germ line cells, from small parts of the basal vasculature or even single vascular cells, a phenomenon known as whole body regeneration (Freeman, 1964; Tiozzo et al., 2008; Voskoboynik et al., 2008). After evisceration, the solitary ascidian *Polycarpa* can replace the entire digestive system and reform a functional branchial sac (Shenkar and Gordon, 2015). Most solitary ascidians can also regenerate the oral and atrial siphons after amputation (Auger et al., 2010; Jeffery, 2015a; Gordon et al., 2019). And uniquely among the chordates, many ascidian species can regenerate the central nervous system, including the neural complex and associated endocrine-like organs (Schultze, 1899; Dahlberg et al., 2009; Medina et al., 2014; Gordon and Shenkar, 2018). As solitary ascidians age, the capacity for regeneration gradually fades and eventually disappears (Dahlberg et al., 2009; Auger et al., 2010; Jeffery, 2012; Gordon and Shenkar, 2018). Although the regeneration capacities of ascidians have been well described, the molecular and cellular mechanisms of regeneration are still incompletely understood (Kassmer et al., 2019).

*Ciona (C. intestinalis* and *C. robusta*) is the most frequently studied solitary ascidian model for regeneration studies (Schultze, 1899; Hirschler, 1911; Jeffery, 2015b). The favorability of this model for regeneration research is due to its relatively large size, body transparency (during youth), and resilience to many different surgical operations. *Ciona* exhibits asymmetric body regeneration, a process in which severed basal body fragments can regenerate missing distal parts, such as the siphons and neural complex, but severed distal fragments are unable to regenerate basal parts, such as the heart, gonads, and other visceral organs, and eventually perish (Hirschler, 1911; Jeffery, 2015b). This type of regeneration contrasts to that of planarians (Reddien and Sanchez-Alvarado, 2004) and *Hydra* (Reddy et al., 2019), in which complete severance of the body leads to the regeneration of both fragments. A requirement for ascidian regeneration seems to be the inclusion of the branchial sac, a massive pharyngeal organ, in a regenerating body fragment (Hirschler, 1911; Jeffery, 2015a). The branchial sac is punctuated by perforations called pharyngeal fissures, which are lined with ciliated cells that propel water through the body cavities for filter feeding and gas exchange (Manni et al., 2002). *Ciona* increase in size rapidly during adult life, largely due to the substantial growth of the branchial sac, which involves the elongation, splitting, and multiplication of the pharyngeal fissures and adjacent tissues (Millar, 1953). Beginning with only two pharyngeal fissures in recently metamorphosed juveniles, mature adult *Ciona* eventually forms a branchial sac consisting of hundreds of pharyngeal fissures aligned in multiple rows perpendicular to the longitudinal axis.

The adult branchial sac also contains multiple transverse vessels with lymph nodes containing niches of adult stem cells (Jeffery, 2015a). The stem cells of the branchial sac divide to produce migratory progenitor cells responsible for replacing distal body parts, including the siphons and neural complex, the repair of wounds, and the replacement of ciliated cells in the pharyngeal fissures, which turnover rapidly during branchial sac growth (Jeffery, 2015b, 2019). A short exposure of regenerating *Ciona* to the cell proliferation marker EdU strongly labels the branchial sac stem cells, and the resulting progenitor cells can be chased into the sites of wounds and regenerating organs (Jeffery, 2015b, 2019). The progenitor and stem cell migrations specifically target the sites of injury or cell replacement, and in cases of multiple injured sites, each of them, but not the uninjured sites, are invaded by cells derived from the branchial sac stem cell niches. During homeostatic growth the same stem cells niches also produce progenitor cells for replenishment of the recycling ciliated cells in the pharyngeal fissures (Jeffery, 2019). Oral siphon regeneration in *Ciona* involves the upregulation of a suite of genes at the site of amputation, including those coding for microRNAs (Spina et al., 2017) and members of the Notch pathway, such as Delta, Notch, and Fringe, cell guidance factors such as netrins, and proteins involved in tissue repair (Hamada et al., 2015). However, the system that activates cell proliferation in the stem cell niches and directs the migration of progenitor cells to their distal body and pharyngeal fissure targets has not been identified.

The diverse sites in the *Ciona* body targeted by branchial sac stem cells have one commonality: they are all regions of extensive and transient apoptotic cell death (Jeffery, 2019). Apoptosis begins at the margin of the excision early after siphon amputation, extirpation of the neural complex, or wounding, and appears to occur continuously during the rapid turnover and replacement of cells lining the pharyngeal fissures during homeostatic growth. Apoptosis is also linked with regeneration in other animals (Bergmann and Steller, 2010), including the regenerating head in *Hydra* (Chera et al., 2009) and the tail in *Xenopus* tadpoles (Tseng et al., 2009), and appears to induce a signaling cascade leading to cell proliferation (Fogarty and Bergmann, 2017). In *Hydra*, a Wnt ligand is produced by dying cells at the wound site, and apoptosis-driven Wnt signaling has been shown to be an integral part of the head regeneration process (Chera et al., 2010). The Wnt signaling pathway has also been shown to be involved in tissue and organ regeneration in other animals (Whyte et al., 2012).

In this investigation, we have employed caspase inhibitors to determine the roles of apoptosis, Wnt signaling, and progenitor cell targeting during *Ciona* asymmetric regeneration and branchial sac homeostasis.

## Materials and Methods

### Animals

*Ciona intestinalis* was collected at Sandwich Harbor near Woods Hole, MA, USA or raised from fertilized eggs at Station Biologique, Roscoff, France. The larvae were allowed to attach and undergo metamorphosis on plastic Petri dishes. The Petri dishes with attached juveniles were placed on racks in aquaria, fed daily with green algae, and small adults were raised to the desired size for operations (1-2 months old with about 8-16 transverse vessels in their branchial sacs) in aquaria with running sea water. For the pharmacological screen, 6-8 cm adults were used, which were farmed in the laboratory from fertilized eggs or freshly collected from the wild.

### Operations

Animals were anesthetized by treatment for 15-20 min with 0.2 mg/mL tricaine methane-sulfonate (MS222; Sigma Aldrich, St. Louis, MO) buffered in Millipore filtered sea water (MFSW). Operations were carried out using straight-bladed micro-cautery scissors or fine dissection scissors (Fine Science Tools, Foster City, CA, USA). Oral siphons were amputated by severing perpendicular to the long axis at a position immediately below the ring of tentacles, as described previously (Auger et al., 2010). Midbody amputations were made through a plane perpendicular to the longitudinal axis at the level of the rectal opening into the atrial cavity. Operated animals were cultured while attached to Petri dishes, which were placed on racks in running sea water aquaria, or were detached and raised for 6-8 days post amputation (PA) in plastic cell wells containing MFSW.

### Caspase inhibition

Animals were treated with 30 μM caspase-1 inhibitor (Calbiochem, San Diego, CA), 30 μM caspase-3 inhibitor (Calbiochem), or 1.5 μM pan-caspase inhibitor Z-VAD-FMK (R & D Systems, Minneapolis, MN). Stock solutions were prepared in dimethyl sulfoxide (DMSO; Sigma Aldrich), frozen, and thawed before each experiment. The effective dose of the caspase inhibitors was determined empirically from the effects of a dilution series on survival and oral siphon regeneration. Caspase inhibitor incubations were done at 16-18°C in the dark.

### Apoptosis detection

Apoptotic cells were detected by terminal deoxynucleotidyl transferase dUTP nick end labeling (TUNEL). To detect apoptosis after oral siphon amputation, amputees were anesthetized at 12 hr PA as described above, their tunics were removed by dissection, the denuded animals were fixed in 4% paraformaldehyde (PFA) for 14 hrs, and washed three times in phosphate buffered saline (PBS), permeabilized in 0.5% Triton X-100, and then washed three more times with PBS. The samples were processed for detection of apoptotic cells with Alexa Fluor azide 488 using the Click-it Plus TUNEL kit (ThermoFisher, Waltham, MA) according to the manufacturer’s instructions, and imaged by fluorescence microscopy. Apoptotic nuclei were quantified in flat mount preparations (Auger et al., 2010) by counting labeled nuclei in 200 μm^2^ areas centered along at the edge of the siphon stump. To detect apoptosis in the pharyngeal fissures of the branchial sac or in the severed ends of distal and basal fragments after midbody amputation, TUNEL was conducted using the In Situ Cell Death Kit (Roche) as described previously (Jeffery, 2019) using animals processed as above. The specimens were post-fixed in 4% PFA in PBS overnight at 4° C, embedded in Paraplast, sectioned at 10 μm, the sections were attached to gelatin subbed glass slides, and unstained sections were imaged by microscopy.

### Carmine assay for branchial sac filtration

Carmine powder (Fisher Scientific, Waltham, MA.) was pulverized with a mortar and pestle. Animals treated with DMSO or caspase inhibitors were mixed with a suspension of 0.1 mg/mL carmine particles in MFSW and incubated for 5 hrs at room temperature. At the end of the assay, animals were photographed using dark field optics.

### Pharyngeal fissure analysis

DMSO and caspase-treated animals were anesthetized as described above and flattened to a thin layer on glass microscope slides by withdrawing most of the MFSW. The pharyngeal fissures were measured along their longitudinal axis using an optic micrometer. Ten fissures were measured along the mid-body row and the values averaged for each animal.

### EdU pulse-chase labeling

Animals treated with DMSO or caspase inhibitors were incubated with 200 μmol/L 5’ ethynyl-2’-deoxyuridine (EdU; Invitrogen, Carlsbad, CA, USA) for 2 days PA at 18°C in MFSW for the EdU pulse. The EdU was chased by five successive washes in MFSW and subsequent culture in MFSW without EdU for 6 days. EdU pulse and chase labeled animals were relaxed by treatment with 2-4 crystals of menthol (Sigma-Aldrich) for 30 mi at room temperature, fixed in 4% PFA for 14 hrs, washed three times in phosphate buffered saline (PBS), permeabilized in 0.5% Triton X-100, washed three more times with PBS, processed for EdU detection with Alexa Fluor azide 488 using the Click-it imagining kit (ThermoFisher), as described previously (Jeffery, 2015a), and imaged by fluorescence microscopy.

### Alkaline phosphatase staining

Animals from the EdU pulse-chase experiments were relaxed in menthol (see above) fixed in 4% formaldehyde for 1 hr at room temperature, washed three times in PBS, and then treated with BCIP-NRH (Invitrogen) for 15-30 min at room temperature in the dark (Jeffery, 2015a). After development of purple color, the animals were washed three times in PBS and imaged.

### Pharmacological screen

The pharmacological screen was carried out using adult animals 6-8 cm in length. The FGF signaling pathway inhibitor was SU5402 (5 μM) (Tocris Bioscience, Bristol, UK) (Grand et al., 2004), the Hedgehog signaling pathway inhibitors were cyclopamine (20 μM) (Chen et al., 2002a) (Tocris Bioscience) and Sant-1 (Tocris Bioscience) (15 μM) (Chen et al., 2002b), the BMP signaling pathway inhibitor was dorsomorphin (Tocris Bioscience) (5 μM), the Notch signaling pathway inhibitors were DAPT (Dovey et al., 2001) (12 μM) (Tocris Bioscience, Bristol, UK) and Compound E (Seiffert et al., 2000) (1 μM) (Abcam, Cambridge, MA, USA), and the Wnt pathway signaling inhibitors were FH535 (1.2 μM) (Santa Cruz Biotechnology Inc., Dallas, TX) (Handeli and Simon, 2008) and IWR-1-Endo (2.5 μM) (Santa Cruz Biotechnology) (Chen et al., 2009). Stock solutions were prepared in DMSO. The effective concentrations were determined by assaying survival and oral siphon regeneration capacity in serial dilutions. Animals were pre-incubated with an inhibitor for 12 h at 16-18°C prior to oral siphon amputation. After oral siphon amputation, the amputees were treated with an inhibitor for 8 days at 16-18°C with fresh changes of inhibitor added every 2 days. The unoperated controls were treated with DMSO in the same final concentration as in the inhibitors.

### Short interfering RNA treatment

Treatment was carried out using siRNAs designed by Invitrogen using NCBI sequence information for *Ciona intestinalis*. The sequence of *ma3* siRNA was 5’-CUUGUUUGAUGAUGUACCUAAGAC-3’, the sequence of *delta1* siRNA was 5’-CCAGUGAAGGCUCUUUCCAAUUGA-3’, the sequence of the scrambled control *delta1* siRNA was 5’-CCAAAGCGGUCUCUUAACUUGUGAA-3’, the sequence of the *ß-catenin* siRNA was 5’-CCAAGUGGUUGUUCAACAATT-3’, the sequence of the scrambled control *ß-catenin* siRNA was 5’-CCAGUUGUUGGUUUACAAGCAATT-3’, the sequence of *wnt3* siRNA was 5’-GCAGUAACGUCCUGGCUAATT-3’, and the sequence of the scrambled control *wnt3* siRNA was 5’-GCAGCAAUCCUCGGUGUAATT-3’. The siRNAs were diluted in RNase-free water to prepare 100 μM stock solutions, which were stored at −20°C prior to use. The effective siRNA concentrations were determined empirically by assaying the effects of a dilution series on survival and oral siphon regeneration. Animals were treated with 2 μM siRNA in MFSW immediately after amputation. At 3 days PA, the animals were rinsed once with MFSW and incubated in fresh 2 μM siRNA, and regeneration was assayed after 6 days of culture at 18°C.

### Wnt3a rescue

Rescue experiments were carried out at 16-18°C in the same way as in the siRNA and caspase inhibitor experiments. Animals were preincubated with 150 ng/mL Wnt3a (R&D Systems) in MFSW for 1 hr prior to amputation, amputated, then incubated with fresh 150 ng/mL Wnt3a after amputation along with an siRNA, caspase inhibitor, or a caspase inhibitor and EdU at 18°C as described above. Fresh Wnt3a was also added to the chase in the EdU labeling experiments. As controls, amputated animals were treated with 150 ng/mL Bovine Serum Albumen (BSA; ThermoFisher), human recombinant BMP-2 (Advent Bio, Elk Grove Village, IL) or human recombinant FGF-10 (Advent Bio) using the same regime as described for Wnt3a.

## Results

### Apoptosis is required for oral siphon regeneration

To determine whether apoptosis is required for regeneration, oral siphons were amputated, and the amputees were immediately treated with caspase 1 inhibitor, caspase 3 inhibitor, pan-caspase inhibitor Z-VAD-FMK, or DMSO, which was used as a control (Fig. 1A-J). At 12 hrs post-amputation (PA), some of the amputated animals were fixed and subjected to TUNEL analysis to determine the effects on apoptosis at the wound sites. At this early stage in the regeneration process (Auger et al., 2010), the wound epidermis had not formed and the oral siphon stumps still showed irregular margins (Fig. 1A-D). In the controls, a band of TUNEL labeling was detected at the amputated margin of the oral siphon (Fig. 1A), as described previously (Jeffery, 2019), but TUNEL labeled cells were significantly reduced in the amputees treated with caspase inhibitors (Fig. 1B-D, I), indicating that apoptosis was suppressed. At 6 days PA, the inhibitor treated and control amputees were assayed for regeneration using the re-growth of the oral siphon, differentiation of new circular muscle bands (CMB) (Jeffery, 2015a), and re-appearance of oral siphon pigment organs (OPO) (Auger et al., 2010) as criteria. Re-growth of the oral siphon, differentiation of multiple rows of new CMBs, and formation of the conventional number of 8 OPO, were seen in the controls (Fig. 1E, J), whereas most of the caspase-inhibitor treated amputees lacked these markers (Fig. 1F-H, J). Apoptosis is a transient step in oral siphon regeneration, which begins shortly after amputation and persists for about 1-day PA (Jeffery, 2019). The pan-caspase inhibitor Z-VAD-FMK was effective in suppressing regeneration when treatment was initiated immediately following oral siphon amputation but not after 1-3 days PA (Fig. 1K), thus linking caspase suppression of regeneration to the period of apoptosis and showing that caspase inhibitors have no effects on regeneration outside the apoptosis period. These results indicate that apoptosis is required for oral siphon regeneration in *Ciona*.

**Figure 1.**
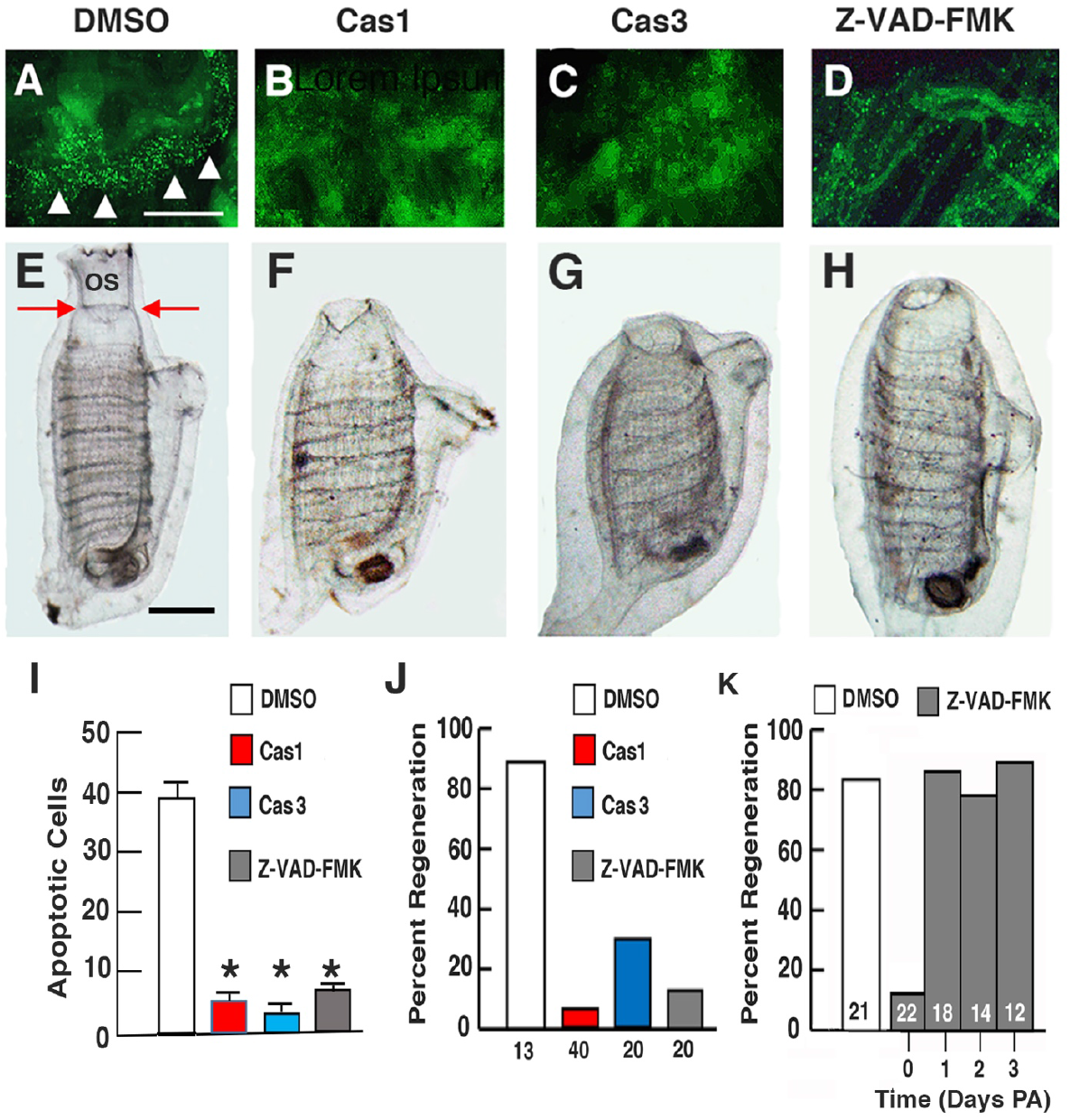
Apoptosis is required for distal body regeneration. A-D. Animals with amputated oral siphons assayed by TUNEL labeling (arrowheads) at the amputation margin after treatment with (A) DMSO (control), (B) caspase 1 inhibitor, (C) caspase 3 inhibitor, or (D) pan-caspase inhibitor Z-VAD-FMK at 12 hrs post amputation (PA). Scale bar: 20 μm; magnification is the same in A-D. E-H. Oral siphon regeneration assayed at 6 days PA after continuous treatment since amputation with (E) DMSO (control), (F) caspase 1 inhibitor, (G) caspase 3 inhibitor, or (H) pan-caspase inhibitor Z-VAD-FMK (H). Arrows in E show position of amputation. OS: oral siphon. Scale bar: 100 μm; magnification is the same in E-H. I. Bar graphs comparing the mean numbers of TUNEL positive nuclei in flat mounted edges of amputated siphon stumps in DMSO, caspase 1 inhibitor-, caspase 3 inhibitor-, and pan-caspase inhibitor Z-DAK-FMK inhibitor-treated animals with amputated oral siphons at 12 hr PA. N = 6 for each bar. Error bars: SEM. Asterisks indicate significant differences at p = > 0.001 between the control and caspase inhibitor treated animals. Statistics by one-way ANOVA and post-hoc Tukey with Bonferroni correction. J. Bar graphs showing the percentage of regeneration at 6 days PA after continuous treatment with DMSO (control), caspase 1 inhibitor, caspase 3 inhibitor, or pan-caspase inhibitor Z-VAD-FMK since the time of amputation. Numbers of animals are indicated at the base or within the bars. K. Bar graphs showing the relationship between the beginning of pan-caspase inhibitor Z-VAD-FMK inhibitor treatment after oral siphon amputation and the percentage of regeneration at 6 days PA. Numbers of animals are indicated within the bars.

### Apoptosis is required for branchial sac homeostasis and function

Apoptosis also occurs during the homeostatic replacement of ciliated cells that line the pharyngeal fissures of the branchial sac (Jeffery, 2019). To determine if apoptosis is required for branchial sac homeostasis, the length of pharyngeal fissures was compared in animals treated with caspase 1, caspase 3, pan-caspase Z-VAD-FMK inhibitors, or DMSO for 6 days (Fig. 2A-E). TUNEL labeling showed that apoptotic cells were abundant in the pharyngeal fissures of the controls (Fig. 2F), but reduced or eliminated in animals treated with the caspase inhibitors (Fig. 2H-I). The pharyngeal fissures were organized in rows and elongated columns typical of growing *Ciona* (Millar, 1953; Manni et al., 2002) in the controls (Fig 2A, E), but the pharyngeal fissures of animals treated with caspase inhibitors were smaller and organized haphazardly (Fig. 2B-E), suggesting that inhibition of apoptosis affected pharyngeal fissure growth and organization.

**Figure 2.**
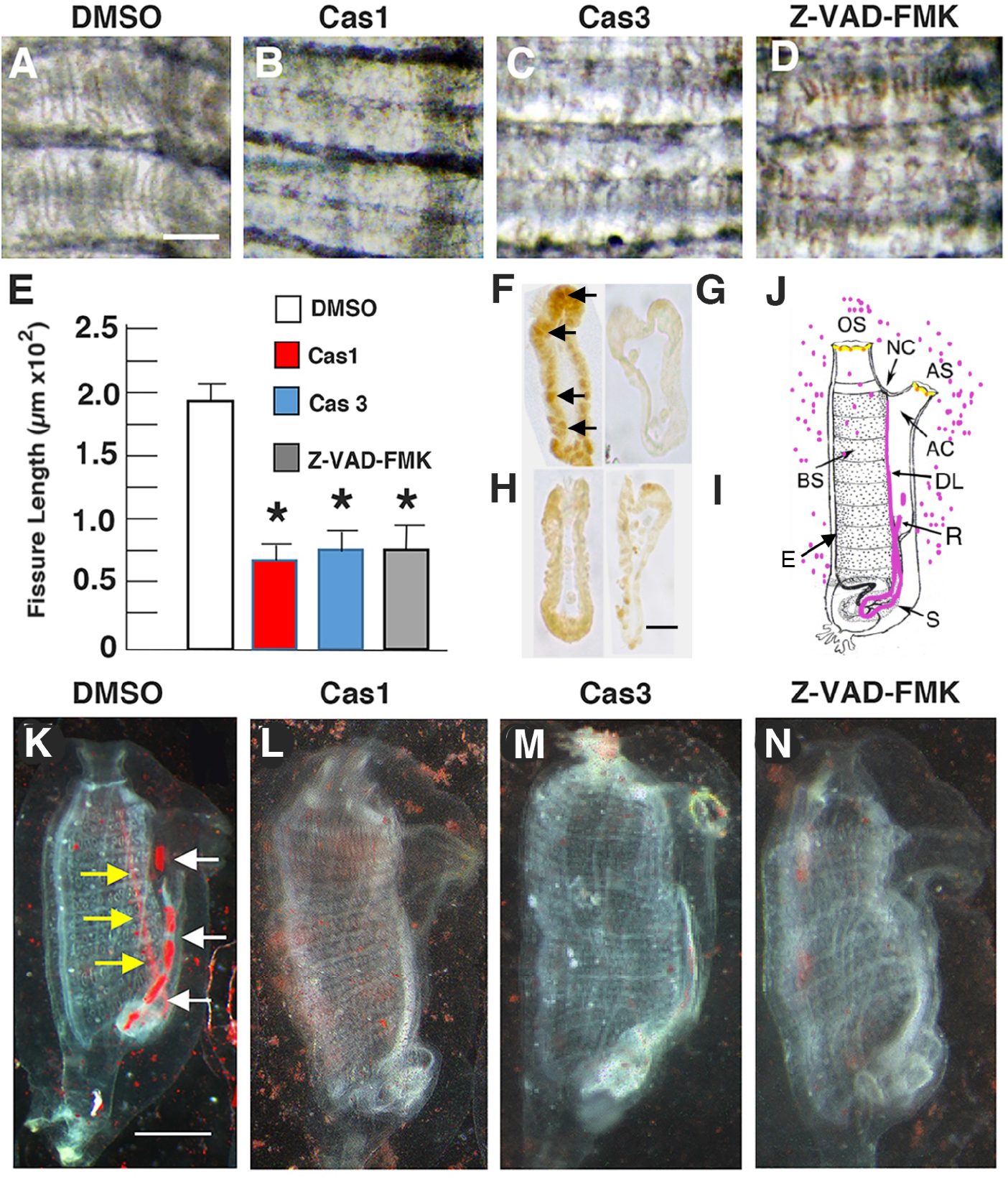
Apoptosis is required for branchial sac homeostasis and function. A-D. Branchial sac fissures after 6-day treatment with (A) DMSO (control) (B) caspase 1 inhibitor, (C), caspase 3 inhibitor, or (D) pan-caspase inhibitor Z-VAD-FMK. Scale bar: 15 μm; magnification is the same in A-D. E. Bar graphs showing the mean lengths of branchial sac fissures in caspase inhibitor and DMSO (control) treated animals. N = 12 for each bar. Error bars: SEM. Asterisks indicate significant difference at p = 0.004 between the control and caspase inhibitor treated animals. Statistics by one-way AVOVA and post-hoc Tukey with Bonferroni correction. F-I. Sections of TUNEL assayed pharyngeal fissures of (F) DMSO (control), (G) caspase 1 inhibitor-, (H) caspase 3 inhibitor, or (I) pan-caspase inhibitor Z-VAD-FMK inhibitor-treated animals. Arrows in F: TUNEL positive nuclei. Scale bar: 5 μM; magnification is the same in all frames. J. Diagram illustrating the carmine particle assay for branchial fissure function. Carmine particles shown by magenta dots and colored organs in the body. OS: oral siphon. AS: atrial siphon. NC: neural complex. BS: branchial sac. DL: dorsal lamina. S: stomach. R: rectum. AC: atrial cavity. K-N. Carmine particle assay of animals treated with (K) DMSO, (L) caspase 1 inhibitor, (M) caspase 3 inhibitor, or (N) pan-caspase inhibitor Z-VAD-FMK. Scale bar: 110 μm; magnification is the same in K-N. Arrows in K show carmine particles concentrated in the dorsal lamina (yellow arrows) and rectum (white arrows).

To determine the effect of caspase inhibitors on branchial sac function, a filtration assay was developed using carmine particles (Fig. 2J). In this experiment, animals were treated with the caspase inhibitors, then incubated in MFSW containing suspended carmine particles, and several hours later the accumulation of carmine particles was examined in the body. During ascidian filtration feeding food particles are trapped in a secretion from the endostyle, which is located on the ventral side of the branchial sac, and collect in the dorsal lamina, which is located on the opposite side of the branchial sac, where a food bolus is formed and passed into the stomach and intestinal tract (Pennachetti, 1984; Petersen and Svane, 2002). The fecal pellets collect in the rectal tube and are ultimately expelled through the atrial siphon (Fig. 2J). The controls showed carmine particles concentrated in the dorsal lamina and rectum (Fig. 2K), indicative of active branchial sac filtration, whereas carmine particles did not accumulate in within the body of animals treated with the caspase inhibitors (Fig. 2L-N), suggesting malfunction of the filtration process. The results suggest that apoptosis in the pharyngeal fissure cells is required for normal growth and function of the branchial sac.

### Role of Wnt signaling in oral siphon regeneration

To explore the molecular basis of oral siphon regeneration, a pharmacological screen was carried out using small molecule inhibitors of classic signaling systems. In these experiments, animals were pre-incubated with an inhibitor, the oral siphon was amputated, incubation with the inhibitor was continued for 6 days, and the extent of regeneration was examined using the markers described above. SU5402 was used to suppress FGF signaling, dorsomorphin to inhibit BMP signaling, and FH535 and IWR-1-Endo to interrupt Wnt signaling. Control amputees were treated with DMSO. In addition, the Notch signaling inhibitors DAPT and Compound E, which were previously shown to suppress oral siphon regeneration in *Ciona* (Hamada et al., 2015), were used as positive controls. As shown in Figure 3A, FGF, Hedgehog, and BMP inhibitors had no effects on oral siphon regeneration. In contrast, the Wnt and Notch inhibitors were effective in suppressing oral siphon regeneration. These results suggest a role for Wnt signaling in oral siphon regeneration.

**Figure 3.**
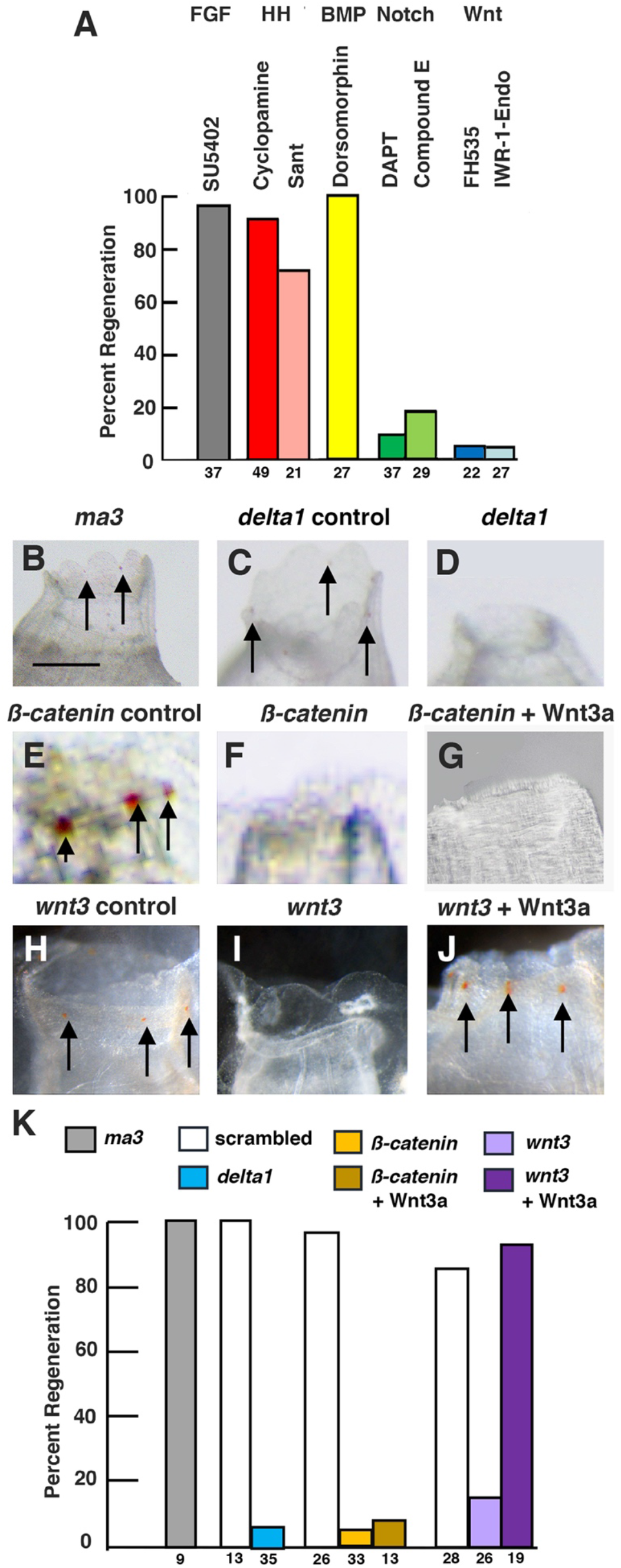
Pharmacological screen and siRNA effects on oral siphon regeneration. A. Bar graphs showing the effects of signaling system inhibitors on oral siphon regeneration. Percent oral siphon regeneration was determined with respect to controls treated with DMSO. The inhibitors and affected signaling systems are shown at the top of the bars. The numbers of animals assayed are shown at the bottom of the bars. B-J. Effects of siRNA on oral siphon regeneration. B, C, E, H. Normal regeneration after treatment with (B) *ma3* siRNA, (C) *delta1* scrambled siRNA, (E) scrambled *ß-catenin* siRNA, or (H) scrambled *wnt3* siRNA. D, F, G. I. Suppression of regeneration after treatment with (D) *delta1* siRNA, (F) *ß-catenin* siRNA, (G) *ß-catenin* siRNA and Wnt3a, or (I) *wnt3* siRNA treatment. J. Rescue of regeneration after treatment with *wnt3* siRNA and Wnt3a. Arrows: oral siphon pigmented organs. K. Bar graphs showing the effect of siRNA on oral siphon regeneration and rescue by Wnt3a. Numbers of animals used in each experiment are shown at the bottom of the bars.

RNA interference is effective for knocking down gene expression in adult ascidians (Rosner et al., 2006; Brown et al, 2009; Tiozzo and De Tomaso, 2009; Rinkevich et al., 2010). Therefore, the role of Wnt in distal regeneration was confirmed using short interfering RNA (siRNA) targeting *wnt3* and *ß-catenin* in the *Ciona* Wnt signaling pathway. *Ciona muscle actin 3* (*ma3*) siRNA was used as a negative control. The *ma3* gene encodes a larval type-muscle actin (Kovilur et al., 1993), which is expressed during embryonic development but not in adults (Chiba et al., 2003), and thus expected to have no effects on regeneration. As a positive control, siRNA corresponding to the Notch pathway gene *delta1*, which is expressed the regenerating oral siphon (Hamada et al., 2015), was used. Lastly, scrambled sequence siRNAs corresponding to the *delta1*, *wnt3*, and *ß-catenin* genes were used as controls. Because *Ciona* constantly filters large volumes of water through the body, an siRNA soaking method was employed in these experiments. Oral siphons were amputated as described above, the amputees were bathed in siRNA, a scrambled sequence siRNA, or DMSO immediately after siphon removal, the original siRNA was exchanged for fresh siRNA at 3 days PA, and regeneration was assayed at 6 days PA using the markers described above. The negative control *ma3* siRNA had no effects on oral siphon regeneration (Fig. 3B, K), and the positive control *delta1* siRNA considerably reduced the numbers of amputees with successful oral siphon regeneration (Fig. 3D, K). The *wnt3* and *ß-catenin* siRNAs also strongly suppressed oral siphon regeneration (Fig. 3F, I, K). None of the scrambled sequence siRNAs had marked effects on regeneration (Fig. 3C, E, H, K). In additional experiments, amputees exposed to *wnt3* or *ß catenin* siRNA were co-treated with human recombinant Wnt3a protein and then assayed for oral siphon regeneration at 6 days PA. The results showed that Wnt3a rescued oral siphon regeneration when applied in combination with *wnt3* siRNA (Fig. 3J, K), but not with *ß catenin* siRNA (Fig. 3G, K), indicating that the effects of *wnt3* siRNA on regeneration are reversible by substitution of a Wnt signaling ligand. Wnt3a, was not expected to restore the effects of ß-catenin siRNA on siphon regeneration because ß-catenin functions downstream in the Wnt pathway. The pharmacological screen and RNA interference experiments support a role for the canonical Wnt pathway in *Ciona* oral siphon regeneration.

### Wnt3a rescues apoptosis inhibition of distal regeneration and branchial sac homeostasis

The role of Wnt signaling in oral siphon regeneration and homeostasis prompted further studies to determine whether exogenous Wnt can rescue the effects of apoptosis inhibition, as has been shown for head regeneration in *Hydra* (Chera et al., 2009).

To determine the effects of Wnt ligand on apoptosis dependent oral siphon regeneration (Fig. 4A), animals were separated into two groups prior to amputation. One group was bathed in Wnt3a protein for 1 hr, and the other group was left untreated. Next, oral siphons were amputated in both groups (Fig. 1E), pan-caspase inhibitor Z-VAD-FMK was added immediately and was present for 6 days PA, after which regeneration was assayed as described above. Most of the amputees treated with pan-caspase inhibitor Z-VAD-FMK did not regenerate amputated oral siphons (Fig. 4B-D), confirming the earlier results (Fig. 1H, J). In contrast, many of the amputees treated with both Wnt3a and pan-caspase inhibitor Z-VAD-FMK showed oral siphon regeneration, including siphon re-growth, CMB differentiation, and OPO formation (Fig 4B, E, F). In contrast, similar exposures of amputees to BSA, FGF, or BMP did not rescue oral siphon regeneration in pan-caspase inhibitor Z-VAD-FMK treated amputees (Fig. 4B). The results indicate that Wnt3a rescues the effects of caspase inhibition on oral siphon regeneration.

**Figure 4.**
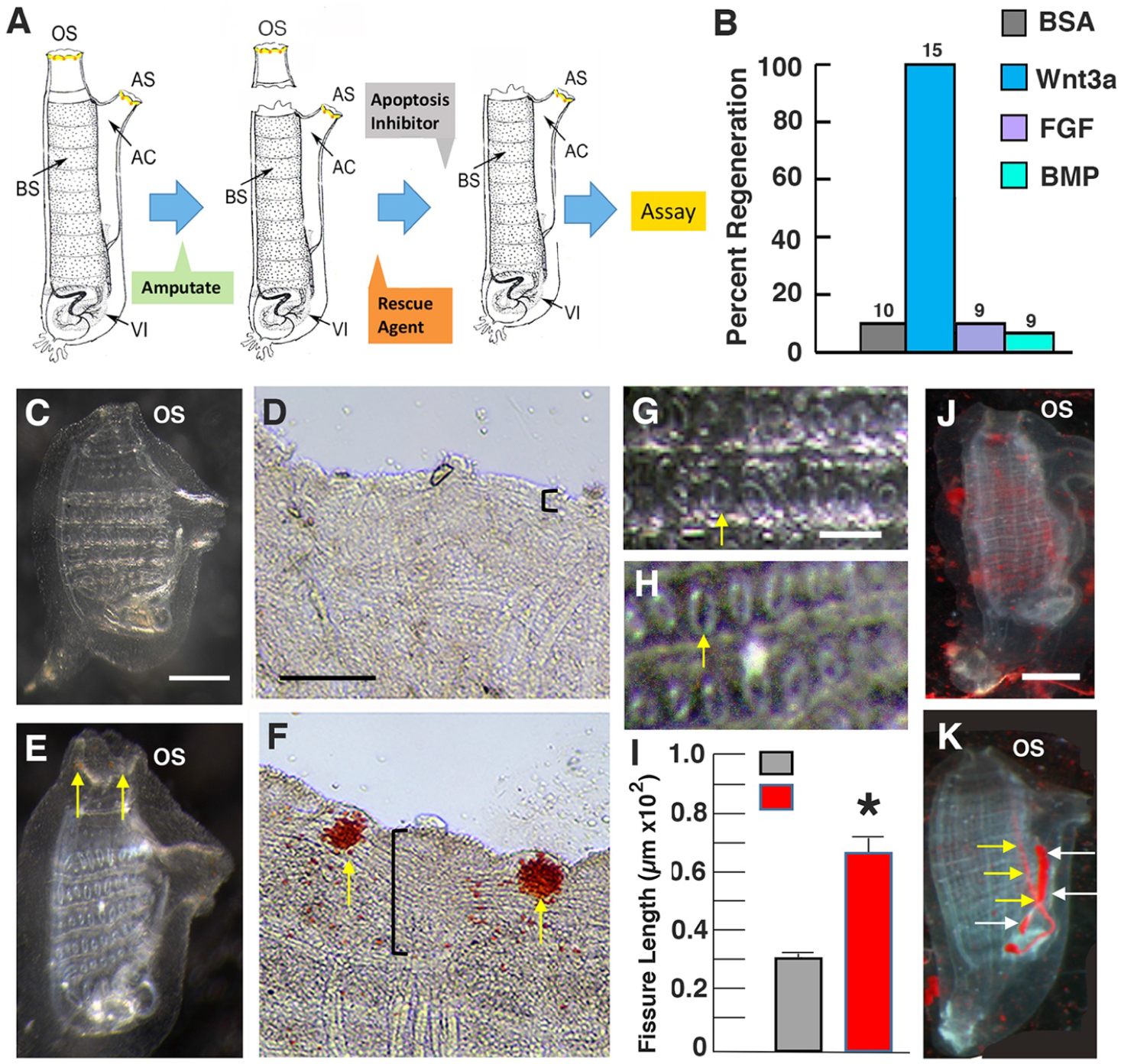
Wnt rescue of apoptosis inhibition effects on oral siphon regeneration and branchial sac homeostasis. A. Diagram of the rescue experiment. Labels are the same as in Figure 2J. B-F. Wnt3a rescue of inhibited oral siphon regeneration. B. Bar graphs showing the percentage of regenerated oral siphons after exposure to pan-caspase inhibitor Z-VAD-FMK and either bovine serum albumen (BSA, control), Wnt3a, FGF, or BMP for 6-days post amputation. The numbers of treated animals are shown at the top of each bar. C-F. Oral siphon regeneration in pan-caspase inhibitor Z-VAD-FMK treated animals without (C, D) or with (E, F) Wnt3a. C, E. Whole animals. D, F. Magnified areas of siphon margin. Arrows: oral siphon pigmented organs. Vertical brackets: extent of circular muscle band replacement. Scale bar in C: 60 μm; magnification is the same in C and D. Scale bar in D: 10 μm; magnification is the same in D and F. G-I. Wnt3a rescue of inhibited pharyngeal fissure growth. G, H. Branchial sacs showing fissure organization in pan-caspase inhibitor Z-VAD-FMK inhibitor treated animals without (G) or with (H) Wnt3a. Arrows: pharyngeal fissures. Scale bar in G: 15 μm; magnification is the same in G and H. I. Bar graphs showing the mean length of pharyngeal fissures (arrows) in pan-caspase inhibitor Z-VAD-FMK treated animals without or with Wnt3a. Error bars: standard deviation. N is 12 for both categories. Asterisk: significance at p = 0.000. Statistical analysis by Student’s t test.

To determine the effects of Wnt ligand on apoptosis-dependent branchial sac homeostasis, animals were separated into two groups, and one group was pre-incubated with Wnt3a for 1 hr and the other group was left untreated, pan-caspase inhibitor Z-DAK-FMK was added to both groups, incubation was continued for 6 days, and the pharyngeal fissures were measured. Treatment with pan-caspase inhibitor Z-VAD-FMK alone reduced the size of pharyngeal fissures, which was reversed by co-incubation with Wnt3a (Fig. 4G, I), suggesting that exogenous Wnt3a rescued the effects of apoptosis inhibition on branchial sac homeostasis.

In summary, the results show that Wnt3a compensates for the negative effects of caspase inhibition on oral siphon regeneration and branchial sac homeostasis, supporting the possibility that apoptosis controls regeneration and branchial sac homeostasis by activating Wnt signaling in *Ciona*.

### Apoptosis is required for progenitor cell survival

Adult stem cells are activated to divide and dispatch progenitors for wound repair and siphon regeneration and are also involved in replacement of recycling pharyngeal fissure cells during homeostatic growth in *Ciona* (Jeffery, 2015a, 2019). To determine the role of apoptosis in stem cell activities, oral siphons were amputated, the amputees were immediately treated with caspase 1 or caspase 3 inhibitor and the cell proliferation marker EdU for 2 days PA. Some animals were then fixed and the EdU pulse was used to determine the effects on progenitor cell formation (Fig. 5B, C), whereas other animals were subjected to a 6-day EdU chase to determine the fate of the EdU labeled cells (Fig. 5E, F). DMSO treated controls were also subjected to the EdU pulse-chase labeling regime (Fig. 5A, D). No differences were found in labeling of progenitor cells in the branchial sac, stem cell niches in controls and caspase-inhibitor treated animals during the EdU pulse (Fig. 5A-C), indicating that apoptosis is not required for the activation of stem cell proliferation. However, striking differences were seen between the controls and caspase inhibitor treated amputees following the EdU chase. In controls, EdU labeled cells were chased into the regenerating siphon and cells of the pharyngeal fissures (Fig. 5D), as described previously (Jeffery, 2015b, 2019). In contrast, EdU labeling was negligible or only a few EdU labeled cells were visible in the caspase inhibitor-treated animals (Fig. 5E, F), implying that most of the EdU labeled cells perished during the chase. Alkaline phosphatase staining (Jeffery, 2015a) of caspase 3-inhibitor treated animals showed that the stem cells were still present after the EdU chase (Fig. 5F, inset). The results suggest that apoptosis is necessary for progenitor cell survival.

**Figure 5.**
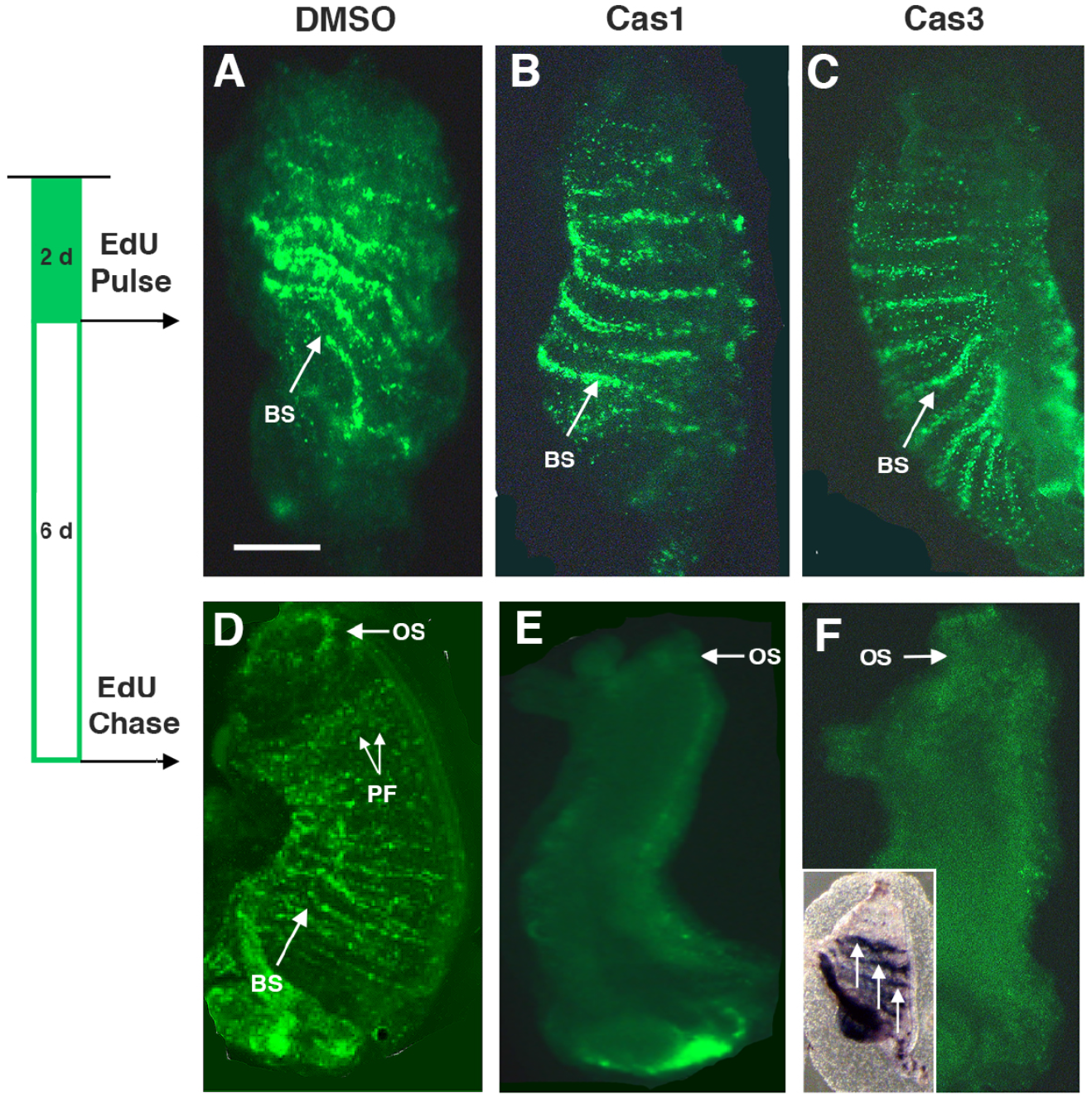
The relationship between apoptosis-dependent oral siphon regeneration, cell proliferation, and survival in the branchial sac stem cell niche. EdU pulse chase labeling regime is shown on the left. Stripes of labeling representing progenitor cells in stem cell niches of the branchial sac after a 2-day EdU pulse (A-C) followed by a 6-day chase (D-F). A, D. DMSO control. B, E. Caspase 1 inhibitor treatment. C, F. Caspase 3 inhibitor treatment. F inset. Alkaline phosphatase staining of stem cell niches (arrows) in the branchial sac of a caspase 3-inhibitor treated animal subjected to the EdU pulse-chase regime. BS: Branchial sac stem cell niche. PF: pharyngeal fissures. OS: oral siphon. Scale bar in A: 100 μm: magnifications are the same in all large frames.

### Apoptosis-dependent Wnt signaling in asymmetric regeneration

In contrast to oral siphon amputation, in which the branchial sac remains unaltered, midbody amputation produces distal and basal body fragments, each containing about one-half of the branchial sac. However, only the basal fragments regenerate (Hirschler; 1914; Jeffery, 2015a; Fig. 6F). Therefore, further experiments were conducted to determine the role of apoptosis-dependent Wnt signaling in asymmetric regeneration.

**Figure 6.**
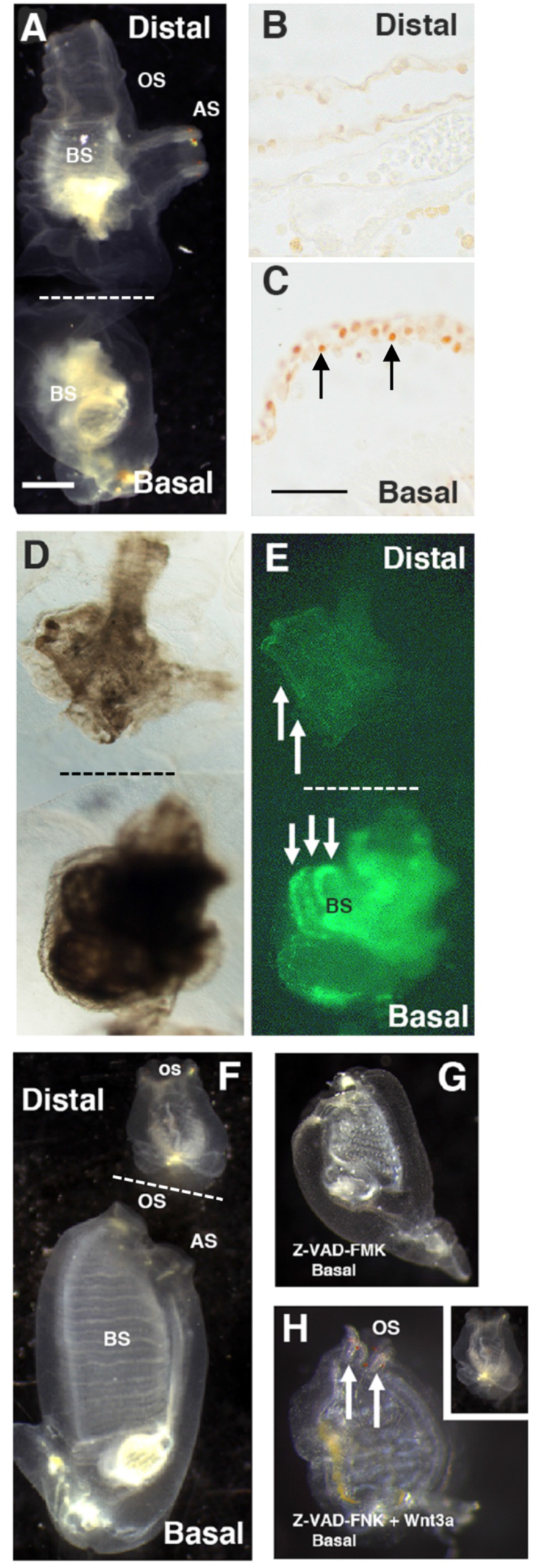
The roles of apoptosis-dependent regeneration and stem cell activation in midbody amputation. A. Bisected animal immediately after mid-body amputation showing distal and contracted basal fragments. Scale bar in A: 100 μm. Magnifications are the same in A and D-H. B, C. Sections of the distal (B) and basal (C) fragments 12 hr after midbody amputation showing TUNEL labeled nuclei (arrows) in the basal but not the distal fragment. Scale bar in C: 10 μm. Magnifications are the same in B and C. D, E. Bright field (D) and fluorescence (E) images of basal and distal fragments subjected to EdU for 2 days after bisection showing progenitor cell labeling in the branchial sac stem cells of the basal (downward arrows) but not the distal fragment (upward arrows). F. Bisected control animal after 6 days post-amputation (PA) showing the regenerating basal fragment and non-regenerating distal fragment. G. A basal fragment treated with pan-caspase inhibitor Z-VAD-FMK immediately after mid-body amputation showing the absence of regeneration at 6 days PA. H. A basal fragment treated with pan-caspase inhibitor Z-VAD-FMK and Wnt3a immediately after mid-body amputation showing rescue of regeneration in the basal fragment (arrows) but not the distal fragment (inset) at 6 days PA. OS: oral siphon. AS: atrial siphon. BS: branchial sac. Dashed lines in A, D-F indicate the bisection plane.

Animals were amputated across the midbody to produce approximately equal-sized distal and basal fragments (Fig. 6A). To address apoptosis, the distal and basal fragments were subjected to TUNEL analysis at 12 hr PA. The severed margin of the basal fragments showed a layer of apoptotic cells (Fig. 6C), but no apoptotic cells were detected at the severed margin of the distal fragments (Fig. 5B), showing that apoptosis is restricted to the basal fragments after mid-body amputation. The severed distal and basal fragments were incubated with EdU for 2 days to determine whether branchial sac stem cells were activated to divide in both fragments after midbody amputation. Progenitor cell labeling was detected in the branchial sac of the basal fragments, but not in the distal fragments (Fig. 6D, E), showing that stem cell activation is unilateral in the basal fragments following mid-body amputation. Midbody-amputated animals were treated with the pan-caspase inhibitor Z-VAD-FMK to determine whether apoptosis is required for basal fragment regeneration. In DMSO controls, the distal fragments showed no regeneration, but the basal fragments had replaced the oral and atrial siphons at 6 days PA (Fig. 6F; Table 1). In contrast, no regeneration was seen in Z-VAD-FMK-treated basal fragments at 6 days PA (Fig. 6G; Table 1), or when observed during the next several weeks, indicating that apoptosis is required for basal fragment regeneration. Siphon regeneration also occurred in basal fragments, but not in distal fragments, treated with both Wnt3a and pan-caspase inhibitor Z-VAD-FM (Fig. 6H and inset; Table 1), which show asymmetric Wnt3a rescue after mid-body amputation. This results indicate that distal fragments are insensitive to Wnt induced rescue of regeneration.

**Table 1.** The effects of pan-caspase inhibitor Z-VAD-FMK and human recombinant Wnt3a on regeneration in distal and basal body fragments after mid-body bisection.

| Treatment<sup>a</sup> | Distal fragments |  |  | Basal fragments |  |  |
| --- | --- | --- | --- | --- | --- | --- |
|  | Regenerated<sup>b</sup> | Total | %Regeneration | Regenerated<sup>c</sup> | Total | %Regeneration |
| DMSO | 0 | 34 | 0 | 19 | 24 | 7 |
| Z-VAD-FMK | 0 | 15 | 0 | 0 | 13 | 0 |
| Z-VAD-FMK and Wnt3a | 0 | 38 | 0 | 10 | 17 | 59 |

- a. Treatments carried as described in Materials and Methods - b. Regeneration determined by appearance of siphons with OPO at severed wound sites on the basal side of the amputated body at 8 days post-operation. - c. Regeneration determined by appearance of siphons with OPO at severed wound site on the distal side of the amputated body at 8 days post-operation.

The results indicate that mid-body amputation shows the same relationship between apoptosis and branchial sac stem cell activation as oral siphon amputation (above and Jeffery, 2015b). Furthermore, the distal fragments of mid-body amputees exhibit no apoptosis at the wound site, no activation of cell proliferation in the branchial sac, and cannot be induced to regenerate by Wnt3a treatment.

## Discussion

This study presents new information on the role of apoptotic cell death at the sites of tissue replacement during regeneration and homeostatic growth in the ascidian *Ciona intestinalis*. We previously showed that apoptosis is an early and transient event in the repair and regenerative programs of wounded animals or amputated distal organs, such as the oral and atrial siphons and the neural complex (Jeffery, 2019). As a consequence of such injuries, adult stem cells in the branchial sac are activated to produce progenitor cells, which migrate through the body and replace injured or missing tissues and organs (Jeffery, 2015). The progenitor cells that replenish blood cells (Ermak, 1976) and rapidly-cycling ciliated cells lining the pharyngeal fissures of the branchial sac (Jeffery, 2019) are also produced by the continuous proliferation of branchial sac stem cells during homeostatic growth. It is shown here that (1) apoptosis is required for distal regenerative activities and normal homeostatic cell replacement in the branchial sac, (2) Wnt signaling is involved in both of these processes, (3) exogenously applied Wnt ligand can substitute for apoptosis, when the latter is inhibited, and rescue normal regenerative and homeostatic activities, and (4) apoptosis is required for the survival, rather than the proliferation, of progenitor cells in the branchial sac stem cell niches. These results suggest an apoptosis-driven Wnt-dependent model for progenitor cell targeting and tissue replacement by adult stem cells during *Ciona* regeneration and homeostasis (Fig. 7).

**Figure 7.**
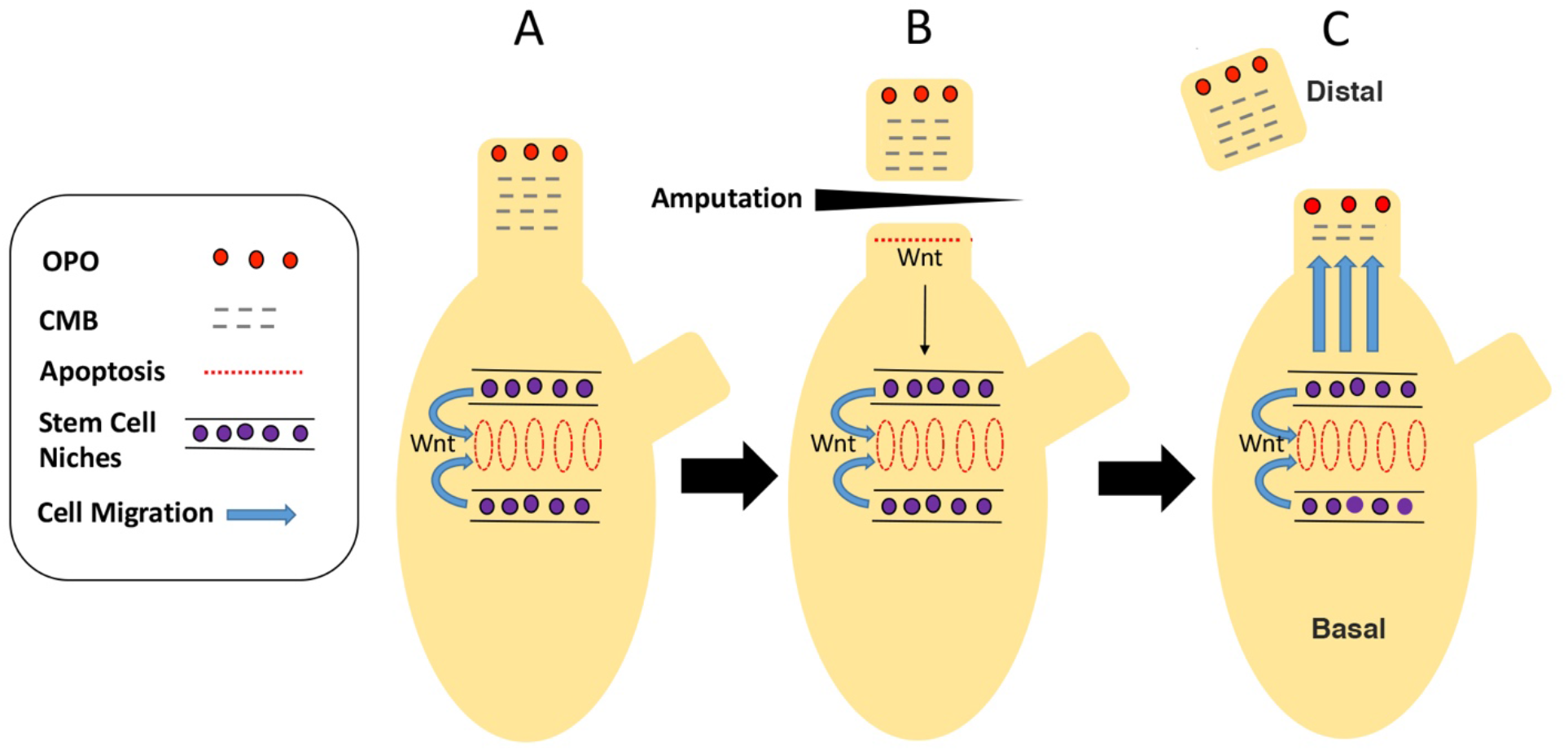
A model for apoptosis-dependent Wnt signaling in oral siphon regeneration and branchial sac homeostasis in *Ciona*. A. Prior to injury, there is no Wnt signaling from the oral siphon stump to the branchial sac stem cell niches. B. Following oral siphon amputation, apoptosis is activated at the proximal margin of the wound, a Wnt signal is generated as a result of apoptosis, and the signal is relayed basally toward the branchial-sac stem cell niches. C. During oral siphon regeneration, Wnt signaling maintains the function of branchial sac stem cells, which target progenitor cells to the oral siphon stump for replacement of lost tissues. A-C. A Wnt signal is continuously generated by apoptotic pharyngeal fissure cells to produce progenitor cells for ciliated cell replacement. OS: oral siphon. AS: atrial siphon. OPO: oral siphon pigment organ. CMB: Circular muscle bands.

The caspase system is essential for the initiation and execution of programmed cell death, and its core components are highly conserved during evolution (Cohen, 1997; McIIwain et al., 2013; Bell and Megeney, 2017). Eleven different caspase genes have been identified in *Ciona*, including genes encoding enzymes responsible for inducing inflammation (e. g. caspase-1) and the initiation and execution of programmed cell death (e. g. caspase-3), suggesting that ascidians utilize a cell death signaling core similar to other animals (Terajima et al., 2003). Our approach was to determine the effects of both specific and general caspase inhibitors on apoptosis associated with *Ciona* regeneration and homeostatic growth. Inhibitors of the inflammatory caspase-1, the apoptotic initiator caspase 3, and the pan-caspase inhibitor Z-VAD-FMK suppressed apoptotic cell death at the apical margin of the amputated siphon stump and prevented the differentiation of new oral siphon tissues and organs, including the CMB and OPO sensory structures. The pan-caspase inhibitor also blocked the regeneration of basal fragments, which show unilateral apoptosis after mid-body bisection. Apoptosis occurs transiently, beginning shortly after oral siphon amputation and continues for about 24 hrs during the beginning of regeneration process (Jeffery, 2019 and unpublished). Caspase inhibition was effective in blocking regeneration only when carried out within the first day after amputation, which spans the period of apoptosis, and inhibitor treatment at later times following amputation had no effects on regeneration. These results indicate that apoptotic cell death at amputation sites is required as an early step in the *Ciona* regeneration program. Early apoptosis is also a critical factor in regenerating planarians (Hwang et al., 2004), *Hydra* (Chera et al., 2009), newts (Vlaskalin et al., 2004), and *Xenopus* (Tseng et al., 2007). The requirement for apoptosis as an early step of regeneration in *Ciona*, an ascidian chordate, provides further evidence for evolutionary conservation of the link between apoptosis and regeneration.

Caspase inhibition also prevented apoptosis in cells lining the pharyngeal fissures of the *Ciona* branchial sac. The ciliated cells are responsible for the flow of food-laden seawater through the body cavities (Millar, 1953; Manni et al. 2002), turnover rapidly during normal growth, and like the progenitor cells responsible for regeneration are derived from branchial sac stem cells (Jeffery, 2019). Caspase inhibition reduced pharyngeal fissure growth and prevented water flow and filtration through the body, consistent with a requirement of apoptosis for recycling of ciliated cells produced by adult stem cells. This process is reminiscent of the role of apoptosis in the recycling, replacement, and growth of adult tissues and organs such as the liver in vertebrates (Fogarty and Bergmann, 2017).

The results suggest that apoptotic cells may initiate a signaling pathway leading to the activation of adult stem cells in the branchial sac and the replacement or replenishment of lost cells and tissues during regeneration and homeostatic growth. The signal could be produced by the dying apoptotic cells themselves or by nearby living cells. Several lines of evidence indicated that canonical Wnt signaling is involved in apoptosis-driven regeneration and homeostasis in *Ciona*. First, regeneration is blocked by small molecule Wnt inhibitors or by siRNA mediated functional inhibition of the *wnt3* and *ß-catenin* genes. The specificity of the latter results was verified in several different ways, most convincingly by the rescue of *wnt3* siRNA effects on regeneration by exogeneous Wnt ligand. Second, exogenously applied Wnt ligand can rescue the effects of caspase inhibitors on regeneration and branchial sac homeostatic growth. Apoptosis-driven regeneration based on Wnt signaling is also involved head regeneration in *Hydra* (Chera et al., 2009), and Wnt signaling has been shown to be central to regeneration throughout the animal kingdom (Whyte et al., 2012). Therefore, our results align *Ciona* regeneration with conclusions obtained from experiments on other regenerating systems. In these situations, it has been shown that apoptosis mediates regeneration by activating the proliferation of stem cells, which has been referred to as a caspase - or an apoptosis-driven proliferation process (Fogarty and Bergmann, 2017). However, according to our results, the situation in *Ciona* may be different. We found that caspase inhibition did not affect the proliferative activity of branchial sac stem cells, but instead appeared to affect the survival (and migration) of progenitor cells derived from the stem cell niches. At least in *Ciona*, the effects of caspase inhibitors on the stem cell niches are thus more accurately described as caspase- or apoptosis-driven cell survival.

Intrinsic differences between oral siphon and mid-body amputation, most importantly the presence of part of the branchial sac and its stem cell niches in distal fragments after the latter operation, led to interesting conclusions about the asymmetric regeneration process at wound sites in *Ciona*. First, apoptosis was unilateral, occurring at the severed margin of the basal but not the distal fragments. Therefore, there seems to be no possibility of apoptosis-driven downstream events in the distal fragments, and this may be a crucial factor in their inability to regenerate. The complete explanation is probably more complex, however, because application of the Wnt ligand, which rescued caspase-blocked regenerative activity in the oral siphon stumps and in the basal fragments produced by mid-body amputation, did not have the same effects on the distal fragments after mid-body regeneration. Second, although the distal fragments of mid-body amputations contain a large part of the original branchial sac, EdU labeling showed that stem cells in this part of the branchial sac did not divide and produce progenitor cells, as occurred in the part of the branchial sac remaining in the basal fragments, and this may be another reason why distal fragments are unable to regenerate. This result also shows that the presence of a branchial sac is not sufficient in itself for regeneration. The first and second conclusions suggest that additional signaling, possibly arising from somewhere in the basal portion of the animal, may also be involved in apoptosis, stem cell proliferation, and regeneration. Future studies with the non-regenerating distal fragments, in particular an assessment of the effects of ectopic apoptosis, may provide further insights into the mechanisms of *Ciona* regeneration.

## Acknowledgements

We thank Gaetan Schires at Station Biologique, Roscoff for *Ciona* culture and Amy Parkhurst and Mandy Ng at the University of Maryland for technical assistance. This work was supported by grants from the National Institutes of Health (AG055411) and EMBRC FRANCE.

## References

Auger, H., Sasakura, Y., Joly, J.-S., Jeffery, W.R. 2010. Regeneration of oral siphon pigment organs in the ascidian *Ciona intestinalis*. Dev. Biol. 339: 374–389.

Bell, R. A. V., Megeney, L. A. (2017). Evolution of caspase-mediated cell death and differentiation: twins separated at birth. Cell Death Differ. 24: 1359–1368.

Bergmann, A., Steller, H. (2010). Apoptosis, stem cells, and tissue regeneration. Sci. Signal 26;3(145):re8.doi: 10.1126/scisignal.3145re8.

Brown, F. D., Tiozzo, S., Roux, M. M., Ishizula, K., Swalla, B. J., De Tomaso, A. W. (2009). Early lineage specification of long-lived germline precursors in the colonial ascidian *Botryllus schlosseri*. Development 136: 3485–3494.

Chen, J. K., Taipale, J., Cooper, M. K., Beachy, P. A. (2002a). Inhibition of Hedgehog signaling by direct binding of cyclopamine to smoothened. Genes Dev. 16: 2743–2748.

Chen, J. K., Taipale, J., Young, K. E., Maiti, T., Beachy, P. A., (2002b). Small molecule inhibitors of Smoothened activity. Proc. Natl. Acad. Sci. USA 99: 14071–14076.

Chen, B., Dodge, M. E., Tang, W., Lu, J., Ma, Z., Fan, C. W., Wei, S., Hao, W., Kilgore, J., Williams, N. S., Roth, M. G., Amatruda, J. F., Chen, C., Lum, L. (2009). Small molecule-mediated disruption of Wnt-dependent signaling in tissue regeneration and cancer. Nat. Chem. Biol. 5:100–107.

Chera, S., Ghila, L., Dobretz, K., Wenger, Y., Bauer, C., Buzgarlu, W., Martinou, J. C., Galliot, B. (2009). Apoptotic cells provide an unexpected source of Wnt3 signaling to drive hydra head regeneration. Dev. Cell 17(2):279–89. doi: 10.1016/j.devcel.2009.07.014

Chiba, S., Awazu, S., Itoh, M., Chin-Bow, S. T., Satoh, N., Satou, Y., Hastings, K. E. M. (2003). A genome wide survey of developmentally relevant genes in *Ciona intestinalis*. IX. Genes for muscle structural proteins. Dev. Genes Evol. 213: 291–302.

Cohen, G. M. (1997). Caspases: the executioners of apoptosis. Biochem. J. 326: 1–16.

Dahlberg, C., Auger, H., Dupont, S., Sasakura, Y., Thorndyke, M., Joly, J-S. (2009). Refining the model system of central nervous system regeneration in *Ciona intestin*alis. PLoS One. 2009; 4(2): e4458. doi: 10.1371/journal.pone.0004458

Dovey, H.F., Anderson, J.P., Chen, L.Z., de Staint Andrieu, P., Fang, L.Y., Freedman, S.B., Folmer, B., Goldbach, E., Holsztynska, E.J., Hu, K.L., Johnson-Wood, K.L., Kennedy, S.L., Kholodenko, D., Knops, J.E., Latimer, L.H., Lee, M., Liao, Z., Lieberburg, I.M., Motter, R.N., Mutter, L.C., Nietz, J., Quinn, K.P., Sacchi, K.L., Seubert, P.A., Shopp, G.M., Thorsett, E.D., Tung, J.S., Wu, J., Yang, S., Yin, C.T., Schenk, D.B., May, P.C., Altstiel, L.D., Bender, M.H., Boggs, L.N., Britten, L.C., Clemens, J.C., Czilli, D.L., Diekman-McGinty, D.K., Droste, J.J., Fuson, K.S., Gitter, B.D., Hyslop, P.A., Johnstone, E.M., Li, W.Y., Little, S.P., Mabry, T.E., Miller, F.D., Audia, J.E., (2001). Functional gamma-secretase inhibitors reduce beta-amyloid peptide levels in brain. J. Neurochem. 76, 173–181.

Ermak, T.H. (1976). The hematogenic tissues of tunicates. In: *Phylogeny of thymus and bone marrow–bursa cells*. Wright RK, Cooper EL (eds), pp. 45–56. Elsevier/North Holland, Amsterdam, The Netherlands.

Fogarty, C. E., Bergmann, A. (2017). Killers creating new life: caspases drive apoptosis-induced proliferation in tissue repair and disease. Cell Death Diff. 24: 1390–1400.

Freeman, G . (1964). The role of blood cells in the process of asexual reproduction in the tunicate *Perophora*. J. Exp. Zool. 156: 157–184.

Gordon, T., Shenkar, N. (2018). Solitary ascidians as model organisms in regenerative biology studies. Results Probl Cell Differ. 65:321–336. doi: 10.1007/978-3-319-92486-1_15

Gordon, T., Manni, L., Shenkar, N. (2019). Regeneration ability in four stolidobranch ascidians: Ecological and evolutionary implications. J. Exp. Mar. Biol. Ecol. 519: https://doi.org/10.1016/j.jembe.2019.151184

Grand E., Chase A., Herath, C., Rahemfulla, A., Cross, N.. C. P. (2004). Targeting FGFR3 in multiple myeloma: inhibition of t (4;14)-positive cells by SU5402 and PD173074. Leukemia 18:962–6.

Hamada, M., Goricki, S., Byerly, M. S., Satoh, N., Jeffery, W. R. (2015). Evolution of the chordate regeneration blastema: Differential gene expression and conserved role of notch signaling during siphon regeneration in the ascidian *Ciona*. Dev. Biol. 405: 304–315.

Handeli, S., Simon, J. A. (2008). A small-molecule inhibitor of Tcf/beta-catenin signaling downregulates PPARgamma and PPARdelta activities. Mol. Cancer. Ther. 7(3):521–9. doi: 10.1158/1535-7163.MCT-07-2063.

Hirschler, J. (1914). Uber die Restitutions-und Involutionsvorange bei operierten Exemplaren von *Ciona intestinalis* Flem. Arch. mikr. Anat., 85, 205–227.

Hwang, J. S., Kobayashi, C., Agata, K., Ikeo, K., Gojobori, T. (2004). Detection of apoptosis during planarian regeneration by expression of apoptosis-related genes and TUNEL assay. Gene 333: 15–25.

Jeffery, W. R. (2001). Determinants of cell and positional fate in ascidian embryos. Int. Rev. Cytol. 203: 3–62.

Jeffery, W. R. (2012). Siphon regeneration capacity is compromised during aging in the ascidian *Ciona intestinalis*. Mech. Ageing Dev. 133: 629–636.

Jeffery, W.R. (2015a). Distal regeneration involves the age-dependent activity of branchial sac stem cells in the ascidian *Ciona intestinalis*. Regeneration 2: 1–18.

Jeffery, W. R. (2015b). Closing the wounds: one hundred and twenty five years of regenerative biology in the ascidian *Ciona intestinalis*. Genesis, doi: 10.1002/dvg.22799.

Jeffery, W. R. (2019). Progenitor targeting by adult stem cells in *Ciona* homeostasis, injury, and regeneration. Dev. Biol. 448: 279–290.

Kassmer, S. H., Nourizadeh, S., De Tomaso, A. W. (2019). Cellular and molecular mechanisms of regeneration in colonial and solitary Ascidians. Dev. Biol. 448: 271–278.

Kovilur, S., Jacobson, J. W., Beach, R. B., Jeffery, W. R., Tomlinson, C. R. (1993). Evolution of the chordate muscle actin gene. J. Mol. Evol. 36: 361–368.

Manni L., Lane N. J., Zaniolo, G., Burighel, P. (2002). Cell reorganization during epithelial fission and perforation: The case of the ascidian branchial fissure. Dev Dyn. 234:303–313.

McIIwain, D. R., Berger, T., Mak, T. W. (2013). Caspase functions in cell death and disease. Cold Spr. Harb. Perspect. Biol. 5(4):a008656. doi: 10.1101/cshperspect.a008656

Medina, B. N., de Abreu, I. S., Cavalcante, L. A., Silva, W. A. B., da Fonseca, R. N., Allodi, S., de Barros, C. M. (2014). 3- acetylpyridine-induced degeneration in the adult ascidian neural complex: reactive and regenerative changes in glia and blood cells. Dev Neurobiol. DOI 10.1002/dneu.22255.

Millar, R.H. (1953). Ciona. In: *LMBC Memoirs on Typical British Marine Plants and Animals*. Colman, JS (ed), pp. 1–123. Liverpool University, Liverpool, UK.

Morizane, A., Doi, D., Kikuchi, T., Nishimjura, K., Takahashi, J. (2011). Small molecule inhibitors of Bone Morphogenic Protein and activin/nodal signals promote highly efficient neural induction from human pluripotent stem cells. J. Neurosci. Res. 89(2):117–26. doi: 10.1002/jnr.22547.

Pennachetti, C. A., (1984). Functional morphology of the branchial basket of *Ascidia paratropa* (Tunicata, Ascidiacea). Zoomorphol. 140: 216–222.

Petersen, J., Svane, I. 2002. Filtration rate in seven Scandinavian ascidians: implications of the morphology of the gill sac. Mar. Biol. 140: 397–402.

Nishida, H. (2002). Specification of developmental fates in ascidian embryos: Molecular approach to maternal determinants and signaling molecules. Int Rev Cytol 217:227–276.

Reddein, P. W., Sanchez-Alvarado, A. (2004). Fundamentals of planarian regeneration. Annu. Rev. Cell Dev. Biol. 20: 725–757.

Reddy, P. C., Gungi, A., Unni, M. (2019). Cellular and molecular mechanism of *Hydra* regeneration. Results Prob. Cell Differ. 68: 259–290.

Rinkevich, Y., Rosner, A., Rabinowitz, C., Lapidot, Z., Moiseeva, E., Rinkevich, B. (2010). Piwi positive cells that line the vasculature epithelium, underlie whole body regeneration in a basal chordate. Dev. Biol. 345: 94–104.

Rosner, A., Paz, G., Rinkevich, B. (2006). Divergent roles of the DEAD-box protein BS-PL10, the urochordate homologue of human DDX3 and DDX3Y proteins, in colony astogeny and ontogeny. Dev. Dyn 235: (6):1508–21. doi: 10.1002/dvdy.20728.

Schultze, L. S. (1899). Die Regeneration des Ganglions von *Ciona intestinalis* L. Jena Z f Naturwiss. 1899; 33: 263–344.

Seiffert, D., Bradley, J.D., Rominger, C.M., Rominger, D.H., Yang, F., Meredith Jr., J.E., Wang, Q., Roach, A.H., Thompson, L.A., Spitz, S.M., Higaki, J.N., Prakash, S.R., Combs, A.P., Copeland, R.A., Arneric, S.P., Hartig, P.R., Robertson, D.W., Cordell, B., Stern, A.M., Olson, R.E., Zaczek, R., 2000. Presenilin-1 and −2 are molecular targets for gamma-secretase inhibitors. J. Biol. Chem. 275, 34086–34091.

Shenkar, N., Gordon, T. (2015). Gut-spilling in chordates: Evisceration in the tropical ascidian *Polycarpa mytiligera*. Sci. Rep. 5: 9614 DOI: 10.1038/srep09614

Spina, E. J., Guzman, E., Zhou, H., Kosik, K. S., Smith, W. C. (2017). A microRNA-mRNA expression network during oral siphon regeneration in *Ciona*. Development 144: 1787–1789.

Terajima, D., Shida, K., Takada, N., Kasuya, A., Roksar, D., Satoh, N., Satake, M., Wang, H-G. (2003). Identification of candidate genes encoding the core components of the cell death machinery in the *Ciona intestinalis* genome. Cell Death Diff. 10: 749–753.

Tiozzo S., Brown, F. D., De Tomaso, A. W. (2008). Stem Cells: From Hydra to Man, Springer, New York, Chapter 6. Regeneration and stem cells in ascidians. Pp. 95–112.

Tiozzo, S., De Tomaso, A. W. (2009). Functional analysis of Pitx during asexual regeneration in a basal chordate. Evol. Dev. 11: 152–162.

Tseng, A-S., Adams, D. S., Qiu, D., Koustubhan, P., Levin. M. (2007). Apoptosis is required during early stages of tail regeneration in *Xenopus laevis*. Dev. Biol. 301: 62–69.

Vlaskalin, T., Wong, C. J., Tslifidis, C. (2004). Growth and apoptosis during larval forelimb development and adult forelimb regeneration in the newt (*Notpthalalmus viridescens*). Dev. Genes Evol. 214: 423–431.

Voskoboynik A, Soen Y, Rinkevich Y, Rosner A, Ueno H, Reshef R, Ishizuka KJ, Palmeri KJ, Moiseeva E, Rinkevich B, Weissman IL. (2008). Identification of the endostyle as a stem cell niche in a colonial chordate. Cell Stem Cell 3:456–464.

Whyte, J. L., Smith, A. A., Helms, J. A. (2012). Wnt signaling and injury repair. Cold Spr. Harb. Prospect. Biol. 4(8):a008078. doi: 10.1101/cshperspect.a008078.

